# A Shared Neural Network for Highly Liked and Disliked Music

**DOI:** 10.1101/2025.01.20.633989

**Authors:** Pablo Ripollés, Amy M. Belfi, Anna Kasdan, Edward A. Vessel, Andrea R. Halpern, Jess Rowland, Robert Hopkins, G. Gabrielle Starr, David Poeppel

## Abstract

While the neural correlates of music-induced peak moments of pleasure (e.g., chills) have been well studied, data on other types of aesthetic responses, such as judgments of *beauty, goodness, or liking,* are sparse. Among these, *liking* is of particular interest given its complex nature and its ubiquity in everyday—and musical—experiences. In this fMRI study, 26 participants listened to musical excerpts while continuously rating how much they *liked* the music. In addition, subjects provided an overall liking judgment at the end of each musical piece. Participants’ individualized musical preferences manifested remarkably early and were sustained over time, with the value of the continuous rating at the end of each piece showing the strongest correlation with the overall ratings. Most importantly, at the neural level, highly liked and disliked musical pieces activated a shared neural network previously associated with the processing of both positive and negative emotional stimuli. This network encompassed the anterior cingulate cortex, limbic system, basal ganglia, and precuneus. Interestingly, participants who reported greater responsiveness to artistic stimuli also showed stronger engagement of this network during music listening overall. Finally, the default-mode network showed more negative modulation by the slope of continuous liking ratings for highly liked than for disliked excerpts, potentially reflecting greater engagement with listening and evaluation as liking increased. In linking behavior and brain function to better characterize the way humans make aesthetic judgments of music, the data support a model in which preferences and aesthetic choices are modulated by brain circuits associated with emotion and reward.

## INTRODUCTION

There is growing interest in studying perception and its neural correlates in the context of complex naturalistic materials, including music [1–18]. Prior work has focused on unveiling the neural correlates of processing various components of music, such as acoustic features [19–22], and components of subjective response, including uncertainty and surprise [23–26], peak moments of pleasure (i.e., musical chills or frissons; [27–33]), or specific emotional experiences [34–36].

But engagment with the arts also involves judgments of beauty, goodness, and liking. A growing body of research examines the brain mechanisms for these responses to music and art, beyond identifying the neural correlates of a work’s perceptual, cognitive, emotional, and reward-related features, [37–47]. These evaluations capture related but distinguishable aspects of aesthetic experience. Here, we focus on music and examine *liking* as a subjective evaluation of how much a listener likes or dislikes a musical excerpt. We consider liking one aspect of aesthetic evaluation, without equating it with judgments of beauty or artistic merit: listeners may like music without judging it beautiful or artistically accomplished. Liking is particularly relevant for music, given its ubiquity in everyday musical experiences and its inherently personal nature: a report of liking expresses the listener’s own response rather than a claim that others are expected to share. In addition, “liking” covers a complex range of experiences. For example, although positive valence is highly correlated with liking [48–50], that is not always the case. We can appreciate art that is unsettling, enjoy horror movies, delight in books or films that end tragically, and, of course, find pleasure in listening to sad songs. Indeed, the emotional valence of music does not need to be in agreement with its aesthetic evaluation (i.e., *sad* music does not always induce sad feelings; note however, that the enjoyment of sad music does not necessarily correspond to a conscious reinterpretation of unpleasant sounds as positive experiences; [51–59]). Examining liking therefore offers a way to investigate how listeners evaluate their musical experiences beyond specific emotional responses; however its neural underpinnings are only partially understood.

Moreover, prior work using naturalistic musical excerpts has typically used (i) listener ratings obtained *post-hoc* or (ii) continuous ratings [60–71]. However, one question that has received less focus is how summary judgments of liking rating relate to judgments as a composition unfolds over time. Real-time measures capture the dynamic fluctuations in emotional or aesthetic responses as the music unfolds, whereas retrospective measures condense that experience into a summary judgment possibly influenced by memory biases (e.g., the peak–end rule; [72]). Studying both provides a more comprehensive understanding of how immediate sensory, cognitive, and affective processes translate into final appraisals, and the neural underpinnings of both. In this vein, various metrics of the continuous response, such as the peak value, slope, or end value, have been used to predict an overall rating, with varying degrees of success [73].

The primary aim of this study was to identify brain networks associated with liking and disliking music. Participants underwent fMRI while listening to musical excerpts and providing continuous liking ratings (i.e., behavioral responses measured dynamically during stimulus presentation; [74]), followed by an overall (post-stimulus) liking judgment after each excerpt. For the musical stimuli, we selected two historically distant and highly distinct genres of music (classical and electronic) to allow for a better generalization across musical style and to provide a range of potentially liked and disliked musical materials. To characterize network-level responsiveness to liking, we used independent component analysis (ICA), a data-driven approach [75] that identifies temporally coherent networks of brain regions. ICA is well-suited to discern how multiple functional networks subserve different cognitive processes [76,77] and has several advantages over univariate analyses (e.g., disentangling different neural circuits occurring within the same brain area; [78–81]).

At the behavioral level, we hypothesized that participants’ liking for the musical excerpts will be apparent very early in the trial, will be sustained over time [62,66], and will show strong agreement with the overall rating after the whole excerpt has been heard. Based on previous research in other aesthetic domains [65,82–86], we also hypothesized low agreement across raters for ratings of individual excerpts (i.e., *one person’s trash is another persońs treasure;* [87–91]).

Most importantly, at the neural level, we hypothesized that aesthetic judgments may show similar neural correlates to prior work in the visual modality (e.g., visual, auditory; [92–95]). Recent studies, which have primarily focused on visual stimuli [38,39,96–98], suggest that aesthetic experiences consistently engage multiple brain networks, including those involved in sensory analysis, emotional responses, and self-referential processes [10,31,35–37,66,96,99–101]. Thus, we hypothesized that networks involved in the processing of music reward and emotion [28,31], as well as self-referential processes [102] would be modulated by continuous and/or overall liking judgments of music.

## MATERIALS AND METHODS

### Participants

Thirty participants were recruited from the greater New York University community. The sample size was selected based on a previous study using a similar behavioral paradigm (i.e., participants providing continuous liking ratings while listening to music; [62]). All participants were right-handed, had no previous history of neurological disorders and reported normal hearing. As prior work has indicated that musical experience as well as expertise influences neural responses to music [103–105], we excluded participants with substantial musical training. Similarly, we excluded participants who were “musically anhedonic,” such that they show no emotional responses to music [68,106–108]. To this end, all participants completed the Goldsmith’s Musical Sophistication Index (Gold-MSI) prior to participating in the study [109]. Participants were excluded if they scored more than 2 SD above the mean on the Musical Training subscale of the Gold-MSI, or more than 2 SD below the mean on the Emotions subscale. Participants were paid for their participation in the experiment. New York University’s Institutional Review Board approved this study, and all participants gave written informed consent in accordance with the New York University Committee on Activities Involving Human Subjects. Three participants experienced technical problems while rating the musical excerpts and one participant did not complete all the fMRI runs after feeling claustrophobic. The reported results are thus based on data from 26 participants (mean age = 27.84 ± 6.21 years; 13 women; Gold-MSI Musical Training Scale = 10.96 ± 3.76, participants’ mean falls within the 11^th^ percentile of the norms; Gold-MSI Emotions Scale = 32.46 ± 4.36, participants’ mean falls within the 31^st^ percentile of the norms; [109]).

### Musical stimulus selection

For classical music, all clips were taken from 19^th^-century small ensemble music from the Romantic era. Because Romantic-era music tends to have a wider range of dynamic and emotional intensity than earlier Classical music, we believed it would allow for a wider range of behavioral responses. No composer was included more than once, and no well-known composers were included; no pieces contained lyrics or the human voice. All the pieces were from the European tradition. Electronic music selections consisted of electronic dance music with a distinctive beat structure (i.e., no ambient music) that contained no lyrics or human vocalizations. All electronic pieces were contemporary and ranged in tempo from 60-150 beats per minute. All stimuli selected for this work have been previously validated and used in a study assessing the timing of music aesthetic judgements [62]. The complete set of stimuli consisted of 16 classical pieces and 16 electronic pieces. Each piece was clipped to a 60s excerpt such that participants could hear enough of the piece to show temporal variability in their ratings, while still allowing for many trials. For electronic music, the 60s excerpts were selected so that some dynamics occurred during that period (e.g., the excerpt contained a “drop” or a transition). Prior to the experiment, participants completed a brief (∼10 minute) training session using musical excerpts from these genres not presented in the experiment to familiarize themselves with the task and response modalities.

### Procedure

The experiment was programmed in PsychToolBox 3.0 [110] using Matlab2017b. Behavioral and fMRI data were collected at the NYU Center for Brain Imaging using a 3T Siemens Allegra scanner. The experiment consisted of four experimental runs with eight musical excerpts each (32 excerpts: 16 classical and 16 electronic). Excerpts were blocked by genre, such that there were two blocks of classical excerpts and two blocks of electronic excerpts. Block and excerpt order were counterbalanced across participants.

In our fMRI experiment, to capture participants’ initial, spontaneous responses to the auditory material, we collected continuous behavioral ratings while participants listened to music, instead of allowing them to listen passively while scanning and collecting liking ratings in a second presentation of the stimuli outside the scanner [17]. Repeated interactions with music can fundamentally alter the experience, including changes in participant engagement and in aesthetic judgments [111–113]. Participants were specifically instructed to use the slider to express how much they liked/disliked the song.

Each trial began with a 1 s blinking fixation cross, followed by the onset of the excerpt. During the excerpt, the participant saw a slider bar presented on the screen. At the onset of music, participants were instructed to continuously rate how much they *liked* the piece at the present moment. To make this continuous rating, participants used a roller ball mouse to move the visual slider bar. The mapping between roller ball and slider was counterbalanced across participants to de-confound movement direction and rating: half of the participants moved the ball up to move the cursor to the right (higher liking) and half the subjects moved the ball down to move the cursor to the right (higher liking). The endpoints of the slider were labeled with ‘L’ and ‘H’ to indicate “Low” and “High”, respectively. Mouse movements were sampled at a rate of 10Hz. Following the offset of the stimulus was a 1s blank screen, which was then followed by an “overall rating” period. During this 4s period, the visual slider bar reappeared on the screen and participants were asked to rate, *overall,* how much they liked the piece. Participants used the slider ball followed by a button-press to make a single, discrete rating on the visual slider bar. For the overall rating, we did not explicitly ask participants to summarize their continuous ratings. Instead, we asked them to report the overall “gestalt” of their experience in terms of liking. Following this rating, the inter-trial-interval (i.e., rest) across all trials was 20s.

After the fMRI experiment had ended, outside the scanner, participants completed the Aesthetic Responsiveness Assessment questionnaire (AReA; [114]). The AReA is a self-report questionnaire that assesses an individual’s general aesthetic responsiveness to stimuli such as paintings, music, natural landscapes, architecture, dance, and poetry, among others. Examples of questions include “I experience joy, serenity, or other positive emotions when looking at art”, “I am emotionally moved by music”, “I appreciate the visual design of buildings”, or “I notice beauty when I look at art”. The English version of AReA has a reliability (ω) of 0.89 and has been validated using several independent datasets and in different languages [114].

Statistical analyses were executed with the software JASP version 0.19.3 [115]. All data were first assessed for normality using the Shapiro-Wilk test. Multiple data distributions deviated from normality and thus non-parametric statistics were used in all analyses (e.g., Wilcoxon signed-rank test), with the matched rank biserial correlation (r_rb_=0.10, small effect size; r_rb_=0.30, medium effect size; r_rb_=0.50, large effect size; r_rb_>=0.7 very large effect size; [116]) or Kendall’s W (W=0.1, small effect size; W=0.3, medium effect size; W>=0.5, large effect size; [117]) as measures of effect size. When needed we computed Bayes factors using default priors (BF_01_), which reflect how likely data are to support the null hypothesis [118–120].

### Behavioral data analyses

#### Inter-rater reliability

To assess the agreement across participants’ liking ratings, we used the intraclass correlation coefficient (ICC). Using JASP, we calculated the “ICC(2,1)” (i.e., a random sample of raters rate all items; [121]) with 95% confidence intervals using the overall ratings from all songs (e.g., 32 songs, 26 raters). An ICC smaller than 0.5 represents ‘poor’ agreement [122].

#### Liking differences between genres

Although we anticipated low inter-rater reliability, it is possible that participants had a general preference for one genre over the other, even if their ratings for individual stimuli within each genre varied widely. To address this possibility, we averaged the overall ratings, and the four measures derived from the continuous ratings for each participant (*end, mean,* and *peak value*, and *slope;* see section below), grouping the values by genre. These average values were analyzed using Wilcoxon signed-rank tests.

#### Relationship between overall and continuous behavioral ratings

First, we tested the relationship between the continuous ratings and the overall rating of liking provided at the end of each stimulus presentation. For each trial, we computed the *slope*, *mean*, *peak* and *end value* as metrics of the continuous trace [73,123]. The slope was calculated using MATLAB r2021b by fitting (in a least-squares sense) a simple linear regression between the liking ratings of each excerpt and the time (i.e., *Y = a + bX*, where *Y* represents the liking ratings, *X* represents time, *a* is the intercept, and *b,* the slope). The slope summarized the direction and average rate of change in liking across the excerpt. Positive slopes indicated an overall increase in liking, negative slopes indicated an overall decrease, and larger absolute slopes reflected steeper changes in ratings over time. The mean value was calculated as the mean of all ratings. For the peak and end value, we first divided the continuous ratings into 2 second bins. Then we averaged the ratings across the bins obtaining 30 values per musical excerpt. We did this to obtain more stable ratings over time. The peak value and the end value were defined as the maximum and the last of these values, respectively. Then, for each participant, we computed Spearmańs correlations between the different metrics of the continuous ratings and the overall responses. We applied Fisher’s z transformation to each participant’s Spearman correlation coefficients. These z-transformed values were analyzed using a non-parametric Friedman test with one factor with four levels (Measure: End Value, Mean Value, Peak Value, Slope). For significant effects, post-hoc Conover tests were used with Holm correction for multiple comparisons. This analysis allowed us to compare the strength of the relationship between the different continuous metrics and the discrete, overall liking score.

For the remaining analyses, trials were designated as ‘high,’ ‘medium,’ and ‘low’ aesthetic appreciation based on the value of the overall rating given at the end of each trial. Specifically, for each participant, the 32 musical excerpts were ranked using these overall ratings and then divided into High (excerpts 1 to 11 in the ranking), Medium (excerpts 12 to 21) and Low (excerpts 22 to 32) trials. This choice of binning was based on previous work showing that neural responses to aesthetic stimuli can follow a nonlinear response based on discrete divisions of aesthetic ratings [96,101]. Thus, binning the responses increased our sensitivity to such a nonlinear response compared to doing an analysis on the continuous response. The remaining behavioral and fMRI analyses were calculated according to this division between High, Medium, and Low.

#### Relationship between acoustic features and behavioral ratings

Previous research has shown that certain acoustic features can be predictive of emotional, aesthetic, and reward-related responses to music [124–127]. We conducted an analysis to test for this possibility. First, we used the MIRToolbox 1.8.1 [128] in MATLAB R2021b to extract acoustic features from each song, employing the default sampling rate and frame lengths. We chose a set of 14 features based on previous studies using music information retrieval techniques [9,125,129–131]. These features represent different aspects of music related to tonality, timbre, temporal properties, and other characteristics [131]. Regarding tonality, we extracted *mode* (using the *mirmode* function; the outcome value represents the difference between the strength of the best fitting major key and the best fitting minor key) and the *key clarity strength* (using the *mirkey* function; the outcome value represents the clarity with which the key of the song can be identified). For timbre, we extracted the *attack time* (using the function *mirattacktime:* the outcome is a vector with the time from the start of an event, identified with *mirevents*, to its peak amplitude; we took the average over all attack times per song), several measures extracted from the spectrum of frequencies (*spectral centroid, spread, flux, flatness,* and *entropy,* using the functions, *mircentroid, mirspread, mirflux, mirflatness, mirentropy,* respectively), and the *roughness* (using the function *mirroughness*; the outcome value represents the sensory dissonance stemming from frequencies with small perceptual distance between them). Regarding temporal features, we calculated the *fluctuation centroid and entropy* (using the *mirfluctuation* function; these measures describe rhythmic periodicities), and the *pulse clarity* (using the *mirpulseclarity* function; the outcome value represents the salience of the song beat). In addition, we also calculated measures related to *loudness* (using the function *mirrms*) and to *novelty* (using the function *mirnovelty,* this measure is related to musical expectations).

For each participant, we then calculated the average values for the 14 acoustic features for the songs in the High and Low liking conditions (i.e., for the top 11 and bottom 11 songs, ranked by the overall liking rating). For each participant and acoustic feature, we thus obtained one average value for songs with high and low ratings. These values were analyzed using Wilcoxon signed-ranked tests.

#### Behavioral Online Replication

To address concerns that scanner noise and physical discomfort might influence the accuracy of participants’ music liking ratings, we conducted an online behavioral replication experiment. This replication followed the same experimental procedures, stimuli, and design as the main fMRI experiment, with the main differences being that: (1) the slider for the ratings went from 1 to 100 instead of 0 to 1 and participants provided responses using their mouse instead of using a roller ball, (2) the experiment was built using oTree [132], and (3) participants were recruited through Amazon Mechanical Turk (AMT). By replicating the experiment in a non-MRI environment, we aimed to validate the consistency and reliability of the behavioral data across contexts. Recruitment was restricted to participants living in the United States and, to ensure data quality, who had completed at least 100 tasks with 95% of approval rate on AMT. Data were collected in summer 2019, before the advent of LLM models. Results from online experiments with participants recruited via AMT up to the widespread use of LLM models have consistently replicated well-established effects [133]. For example, studies have demonstrated that AMT samples yield results comparable to those from traditional participant populations across various cognitive tasks, including, for example, flanker tasks [133,134], experiments involving complex auditory stimuli [135], and experimental designs that require participants to provide continuous ratings while listening to music [60,64]. To address the higher attrition rates typically observed in online experiments [136], we recruited 50 participants. At the end of the experiment, participants completed the Musical Training Scale from the Gold-MSI [109]. In this replication, we did not exclude participants with substantial musical training. To better control for participant characteristics in the online setting and exclude individuals with specific music anhedonia, we administered the Barcelona Music Reward Questionnaire (BMRQ) and excluded participants with scores below 63 [108]. The online experiment was approved by the Institutional Review Board at New York University. Participants provided informed consent by clicking an “accept” button after reading the consent form and were compensated for their participation. Eight participants were excluded from the analysis (1 had specific music anhedonia and 7 did not provide continuous ratings while listening to music). The reported results are thus based on data from 42 participants (mean age = 39.09 ± 12.13 years; 21 women; Gold-MSI Musical Training Scale = 24.76 ± 10.59, participants’ mean falls within the 43^rd^ percentile of the norms; BMRQ=81.28 ± 8.84; for BMRQ scores, a score in the 10th percentile is about 65 and a score in the 90th percentile is about 87; [108,109]).

### fMRI data analysis

#### Scanning parameters

All fMRI scans took place at the NYU Center for Brain Imaging using a 3T Siemens Allegra scanner with a Nova Medical head coil (NM011 head transmit coil). For the music fMRI task, four runs of 346 sequential whole-brain multi-echo (ME) echo-planar imaging (EPI) volumes were acquired (TR = 2000 ms, right-to-left phase encoding, flip angle = 75°, voxel size = 3.0 × 3.0 × 3.0 mm^3^, 34 axial slices, acquisition size = 80×64). The ME EPI sequence and a tilted slice prescription (15-20° tilt relative to the AC-PC line) were used to minimize dropout near the orbital sinuses. ME EPI images were reconstructed using a custom algorithm designed by the NYU Center for Brain Imaging to minimize dropout and distortion. In some of the participants, the sequence did not provide full coverage of the whole brain anatomy, and the most superior parts of the brain were left out (3 mm of the most superior part of the SMA and parietal cortex). To allow precise co-registration with the functional data, a high resolution T1 MPRAGE image was also acquired during this MRI session (TR = 2500 ms, TE = 3.93 ms, flip angle = 8°, voxel size = 1.0 × 1.0 × 1.0 mm^3^, 176 sagittal slices, acquisition matrix = 256 × 256).

#### Preprocessing and ICA analysis

Data were preprocessed using Statistical Parameter Mapping software (SPM12, Wellcome Trust Centre for Neuroimaging, University College, London, UK, www.fil.ion.ucl.ac.uk/spm/) under MATLAB r2021b. For each participant, the four functional runs were first realigned to the mean image of all the EPIs. The high resolution T1 was then co-registered to this mean functional image and segmented using the Unified Segmentation algorithm [137]. The deformation fields (i.e., normalization parameters) obtained during the segmentation process were then used to spatially normalize all functional images to the MNI template included with SPM12 (maintaining the voxel size of 3.0 × 3.0 × 3.0 mm^3^). Finally, images were spatially smoothed with an 8 mm FWHM kernel.

Once fMRI data were preprocessed, Group Spatial ICA was applied using the Group ICA of fMRI Toolbox (GIFT v4.0b; http://mialab.mrn.org/software/gift/; [76]). The number of independent components (formed by a spatial map and a time-course) to be extracted was set to twenty, which has been shown to be an optimal dimension in previous research in both healthy and clinical populations [138–141]. Data were intensity normalized, and a principal component analysis (PCA) was first applied to each participant’s data. The reduced data were then concatenated across participants along the time dimension and reduced again with group-level PCA to 20 dimensions. Infomax ICA [142] was then applied to estimate 20 group components. No scaling was used, as with the intensity normalization step, the intensities of the spatial maps were in percentage of signal change. To obtain whole brain group-wise statistics, the spatial map of each of the ICA components (i.e., networks) retrieved for all participants was submitted to a second-level analysis using a one sample t-test under SPM12, which treats each subject’s spatial map as a random effect [76]. For the spatial maps of each ICA network, all statistics are reported at a p < 0.05 FWE-corrected threshold at the cluster level, with a p < 0.001 uncorrected threshold at the voxel level. Clusters with fewer than 50 voxels were not included in the analyses and are not depicted in the figures. Maxima and all coordinates are reported in MNI space. Anatomical and cytoarchitectonical areas were identified using the Automated Anatomical Labeling [143] and the Talairach Daemon database atlases [144] included in the xjView toolbox (http://www.alivelearn.net/xjview/).

#### ICA Network modulation by music, and by overall and continuous liking ratings

To identify which of the networks were related to the task (i.e., music listening) and to the overall or continuous liking ratings of interest, several multiple regressions were conducted. GIFT allows for fitting each participant’s component time-course to a specific model. The models can be created using SPM12, by convolving a canonical hemodynamic response with the timing of specific conditions and their parametric modulators. First, we visually inspected each network, and those reflecting artifacts such as movements, ventricles, edges, or the presence of blood vessels were discarded. For the non-artifactual components, and to discard networks not modulated by music, we created a model including only one main condition of interest: music (including all musical excerpts regardless of their liking rating). A resting condition plus six movement regressors (obtained from the realignment step, added to take into account motion artifacts) were also included in the model. A multiple regression between this model and each network time-course was fitted and beta values—which represent network engagement—for the music condition were obtained (i.e., we obtained one beta value per network and participant). These models were fitted separately to each participant’s network time courses. These beta values were further analyzed using Wilcoxon signed-ranked tests to identify networks which were significantly activated or deactivated by music listening. In order to control for the number of networks being tested, we adjusted the p-values using FDR-correction as implemented in the *mafdr* function on MATLAB r2021b, which follows the Benjamini and Hochberg method [145].

Once the networks significantly engaged during music listening were identified, a series of finer-grained analyses were performed using the High, Medium, and Low liking categories defined separately for each participant, so that our analysis was more sensitive to a nonlinear response (see Materials and Methods). This binning was done separately for each individual participant, as participants tend to show very low agreement in terms of which pieces are rated ‘high’ versus ‘low’ (see Figure 1A for examples of this disagreement and ICC Results section). Therefore, the same piece may appear in each of the ‘high,’ ‘medium,’ and ‘low’ conditions depending on the participant’s preferences. We compared the acoustic properties of highly liked and disliked songs (see Table 2); however, we found no significant differences for any of the 14 acoustic features tested.

**Figure 1.**
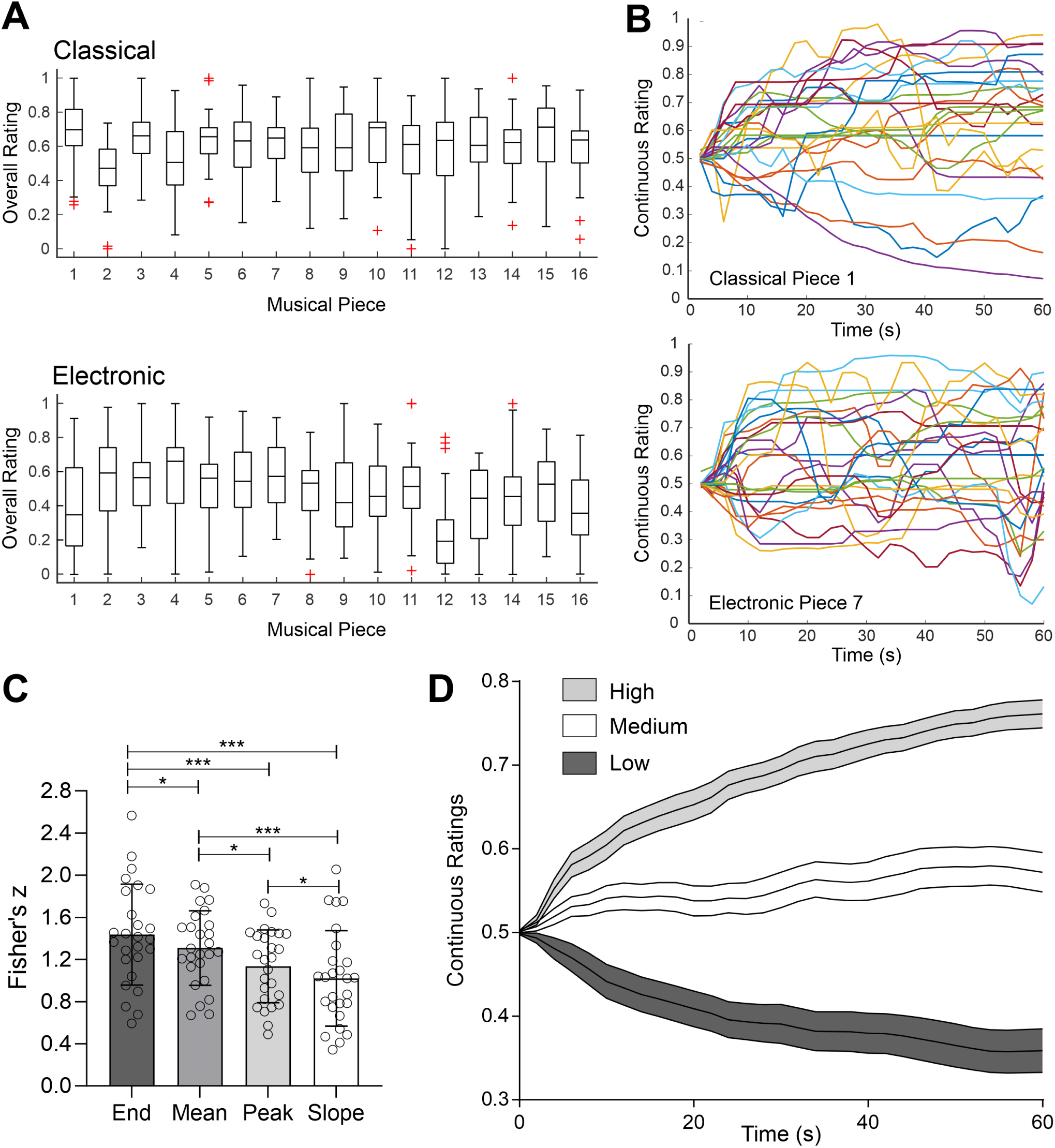
Continuous and overall behavioral aesthetic judgments. **A.** Boxplots illustrating participants’ overall ratings for each musical piece. Center line indicates median, bottom and top edges of the box indicate 25^th^ and 75^th^ percentiles. Whiskers extend to the most extreme data points not considered outliers, and red + symbols indicate individual outliers. **B.** Individual continuous responses from all participants for two representative musical excerpts (top, classical piece number 1; bottom, electronic piece number 7). Each line represents the continuous rating provided by a specific participant. **C**. Barplots (mean ± SD) showing the strength (using Fisher’s z transform) of the correlation between the *overall* rating and different measures extracted from the continuous ratings, including the *end, mean,* and *peak value*, and the *slope.* Each circle represents data from one participant. * p < 0.05, *** p < 0.001. **D.** Mean (solid lines) plus SEM (shaded error bands) of *continuous* ratings (between 0 and 1) across participants are shown for High (grey), Medium (white) and Low (black) musical excerpts classified using the overall ratings.

Once songs were classified into High, Medium, and Low liking conditions, we fitted two models to the networks significantly engaged by music. While both models included these three conditions plus the rest condition and the six movement regressors, they differed in their parametric modulators: one model used the *overall* judgments of liking and another the *slope* of the continuous ratings. The condition regressors and parametric modulators served different purposes. The High, Medium, and Low regressors accounted for the average network response in each category, whereas the parametric modulators tested whether activity varied with ratings or rating slopes within that category.

The first model was used to measure which networks were modulated by the overall liking judgments. A single parametric modulator per musical condition (one for High, Medium and Low trials) was included in this model. This modulator was formed by the ratings provided by participants after the end of each musical excerpt for each condition (High, Medium, Low). The second model was used to identify which networks were modulated by the continuous liking ratings. For each trial, we fitted a linear model to the continuous ratings as a function of time and extracted its slope. Positive slopes indicated that liking generally increased as the music unfolded, whereas negative slopes indicated that it generally decreased. Larger absolute slopes represented steeper changes in ratings over time, while slopes near zero indicated little overall linear change. We used the slope as a trial-level summary of the overall direction and rate of change in liking across each excerpt for two reasons. First, participants’ ratings typically shifted early and thereafter only fluctuated within a relatively stable range (Figure 1B), motivating our focus on the overall trend rather than individual fluctuations. Second, the slope was less strongly associated with overall liking ratings than were the end, mean, or peak values, suggesting that it captures a partly different aspect of the behavioral response (see Figure 1C). A separate slope-based parametric modulator was included for each of the High, Medium, and Low liking conditions.

A multiple regression between these two models and each network time-course was fitted and beta values for each parametric modulator were obtained. To normalize for individual differences, the beta values for the parametric modulator of the Music Medium condition were subtracted from the beta values of the parametric modulators of the Music High and the Music Low conditions for each participant. Thus, for each network, we obtained a series of corrected (from now on, High_m_ and Low_m_) beta values for a) the overall parametric modulator, and b) the continuous parametric modulator (i.e., slopes). The rationale behind this correction by the Medium condition is twofold, based on: (1) the fact that people show great variability in the way in which they process aesthetic stimuli [82,83,85,86]; and (2) the fact that variability in processing aesthetics has direct effects on brain activity within reward and emotion-related regions [30,146–148]. Thus, in this analysis, the Medium condition becomes an intrinsic baseline. We then focused on how high and low aesthetic stimuli *displace* this baseline in opposite or equal directions [149,150]. Finally, for each network and condition of interest (continuous and overall parametric modulators) we used Wilcoxon signed-ranked tests with FDR correction to assess whether the network was significantly modulated by: (1) High_m_ trials; 2) Low_m_ trials; or (3) significantly more modulated by High_m_ than by Low_m_ trials.

## RESULTS

### Behavioral Results

#### Inter-rater variability

Replicating prior work in other aesthetic domains (poetry; visual arts; [82,85,86,151,152]), ICC for overall liking responses was quite low, reflecting disagreement among raters (ICC value= 0.069, 95% CI: 0.033, 0.138). See Figure 1A and 1B for a visual depiction of the variability of the ratings. This low agreement supports the choice of our materials: we wanted pieces/genres on which people would disagree, to reduce reactions to low-level acoustic features.

#### Liking differences between genres

Even in the context of low inter-rater agreement, participants consistently liked classical music more than electronic music for all measures, including both overall and continuous ratings (see Table 1, top).

**Table 1.** Liking ratings for classical and electronic music for the in-scanner experiment (top) and the online behavioral replication (bottom). Ratings are mean ± SD on the original scales (0–1 in the scanner and 1–100 online). Slopes are mean rating changes per second, expressed in points on a 100-point scale for both experiments. *W* is the Wilcoxon signed-rank test, and *r*_rb_ is the matched rank-biserial correlation.

| **Measure** | **Classical** | **Electronic** | ***W*** | ***p*** | ***r*<sub>rb</sub>** |
|---|---|---|---|---|---|
| **In-scanner experiment** |  |  |  |  |  |
| Overall liking | 0.59 ± 0.12 | 0.47 ± 0.10 | 49 | < 0.001 | 0.72 |
| End value | 0.62 ± 0.13 | 0.50 ± 0.12 | 50 | < 0.001 | 0.71 |
| Mean value | 0.59 ± 0.10 | 0.49 ± 0.10 | 54 | 0.001 | 0.69 |
| Peak value | 0.71 ± 0.10 | 0.65 ± 0.09 | 65 | 0.004 | 0.63 |
| Slope | 0.40 | 0.006 | 50 | < 0.001 | 0.71 |
| **Online behavioral replication** |  |  |  |  |  |
| Overall liking | 60 ± 18 | 49 ± 14 | 251 | 0.011 | 0.44 |
| End value | 62 ± 19 | 50 ± 14 | 250 | 0.011 | 0.44 |
| Mean value | 58 ± 14 | 50 ± 10 | 248 | 0.010 | 0.45 |
| Peak value | 73 ± 13 | 66 ± 9 | 229 | 0.005 | 0.49 |
| Slope | 0.29 | 0.024 | 271 | 0.023 | 0.40 |

#### Relationship between overall and continuous behavioral ratings

There was strong agreement between the overall ratings and various metrics extracted from the continuous responses (see Figure 1C). However, there was a significant effect [X^2^ (3) = 38.35, p < 0.001, Kendall’s W = 0.49] with the *end value* of the continuous judgment (see Figure 1C) showing the strongest correlation with the overall ratings, significantly higher than those obtained using the *mean value* (p_holm_ = 0.030, r_rb_ = 0.57), the *peak value* (p_holm_ < 0.001, r_rb_ = 0.81), and the *slope* (p_holm_ < 0.001, r_rb_ = 1).

The *mean* value showed a significantly stronger correlation with the overall ratings than the *peak* value (p_holm_ = 0.013, r_rb_ = 0.67) and the *slope* (p_holm_ < 0.001, r_rb_ = 0.79). The *peak value* also showed a stronger relationship with the overall ratings than the *slope* (p_holm_ = 0.013, r_rb_ = 0.36). This suggests that there was a strong alignment between the overall and continuous ratings, with the *slope* being the metric showing the weakest relationship. As described in the methods, for the remaining analyses we divided the trials according to the overall rating bin (high, medium, low). As expected, the evolution of the behavioral continuous ratings revealed clear differences between high, medium and low-rated musical excerpts (Figure 1D). Strikingly, participants’ average preferences for the musical excerpts were shown very early in the trial (within the first 3 seconds) and were sustained over time.

#### Relationship between acoustic features and behavioral ratings

For each participant in the fMRI experiment, acoustic features were averaged for high and low rated songs. For the 14 features extracted, no significant differences were found between the acoustic characteristics of these songs (see Table 2).

**Table 2.** Acoustic Features. Comparison of average acoustic features for highly liked vs. disliked songs for each participant.

| **Feature** | **W** | **p-value** | ***r<sub>rb</sub>*** |
|---|---|---|---|
| Tonality: Mode | 173 | 0.960 | -0.014 |
| Tonality: Key Clarity Strength | 154 | 0.600 | -0.123 |
| Timbre: Attack Time | 229 | 0.181 | 0.305 |
| Timbre: Spectral Centroid | 124 | 0.199 | -0.293 |
| Timbre: Spectral Spread | 123 | 0.190 | -0.299 |
| Timbre: Spectral Flux | 123 | 0.190 | -0.299 |
| Timbre: Spectral Flatness | 119 | 0.157 | -0.322 |
| Timbre: Spectral Entropy | 110 | 0.099 | -0.373 |
| Timbre: Roughness | 139 | 0.367 | -0.208 |
| Temporal: Fluctuation Centroid | 224 | 0.227 | 0.276 |
| Temporal: Fluctuation Entropy | 213 | 0.353 | 0.214 |
| Temporal: Pulse Clarity | 107 | 0.084 | -0.390 |
| Other: Loudness | 132 | 0.280 | -0.248 |
| Other: Novelty | 201 | 0.532 | 0.145 |

#### Behavioral online replication

The online behavioral experiment replicated the behavioral results obtained for the main fMRI experiment: (1) we found very low agreement among overall liking ratings (ICC value= 0.092, 95% CI: 0.054, 0.163); (2) participants consistently liked classical music more than electronic (see Table 1, bottom); (3) regarding the relationship between overall and continuous ratings, there was a significant effect [X^2^ (3) = 54.88, p < 0.001, Kendall’s W = 0.44] with the *end value* of the continuous judgment showing the strongest correlation with the overall ratings, significantly higher than with any other measure [*mean value*: p_holm_ < 0.001, r_rb_ = 0.60; *peak value:* p_holm_ < 0.001, r_rb_ = 0.81; *slope*: p_holm_ < 0.001, r_rb_ = 1]; and (4) when binning trials into high, medium, and low liking according to the overall ratings, participants’ average preferences for the musical excerpts were shown very early in the trial (within the first 3 seconds) and were sustained over time. See Supplementary Figure 1 for a summary of these results.

### fMRI Results

#### ICA decomposition and network modulation by music

From the original 20 networks extracted using the Group ICA of fMRI Toolbox (GIFT; see Materials and Methods), 6 components reflecting artifacts such as noise, movements, ventricles, edges, or the presence of blood vessels were discarded. Of the 14 remaining components, 11 were significantly and robustly modulated by music listening (by fMRI blocks in which music was presented). While 20 networks have been shown to be an optimal dimension for ICA analysis in previous research in both healthy and clinical populations [138–141], we repeated the analysis extracting 25 networks. The results were very similar, with the same networks being modulated by music listening. Note that most of these networks are typically found in task-related and resting state ICA analyses [141,153–155].

Of these networks, six were positively modulated by music (see Table 3): an *Auditory-Premotor Network* (APMN) covering the bilateral superior and middle temporal gyrus, and premotor and supplementary motor regions; a *Ventromedial Prefrontal Network* (VMPN), covering the ventromedial prefrontal cortex and extending to the ventral striatum and caudate; a network covering the anterior cingulate cortex and frontal regions, and extending to the limbic system, the basal ganglia, posterior cingulate cortex and precuneus, which closely resembles a network previously known to respond to stimuli with both positive and negative emotional valence (a *Valence Network*, VN; [156,157]); an *Orbitofrontal Network* (OrbN), encompassing the lateral and medial orbitofrontal cortex and regions in the occipital cortex; a *Superior Frontal Network* (SFN), extending from superior frontal regions to angular and temporal gyri; and a *Medial Temporal Lobe Network* (MTLN) encompassing the hippocampus, parahippocampus, amygdala and fusiform gyrus.

**Table 3.** ICA networks *positively* modulated by music listening. The main areas of the network are reported using a p < 0.05 FWE-corrected threshold at the cluster level. The level of network engagement with the task is reported using Wilcoxon signed-rank tests and FDR-corrected p-values. APMN: Auditory-Premotor Network; VMPN: Ventromedial Prefrontal Network; VN: Valence Network; OrbN: Orbitofrontal Network; SFN, Superior Frontal Network; MTLN, Medial Temporal Lobe Network.

| **Network** | **Activation Region** | ***W*** | ***p<sub>FDR</sub>*** | ***r<sub>rb</sub>*** |
|---|---|---|---|---|
| **APMN** | Bilateral sup/mid temporal gyrus; Bilateral Heschl gyrus; Bilateral insula; Right precentral gyrus; Bilateral postcentral gyrus; Supplementary motor area. | 351 | <0.001 | 1 |
| **VMPN** | Ventromedial prefrontal cortex; Bilateral anterior cingulate; Bilateral ventral striatum; Bilateral caudate. | 342 | <0.001 | 0.949 |
| **VN** | Bilateral ant/mid/superior cingulate cortex; Bilateral sup/inf frontal gyrus; Bilateral orbitofrontal cortex; Bilateral insula; Bilateral caudate; Bilateral putamen; Bilateral middle temporal gyrus; Bilateral hippocampus; Bilateral parahippocampus; Bilateral amygdala; Thalamus; Bilateral precuneus; Bilateral calcarine; Bilateral lingual gyrus; Right mid/inf occipital gyrus. | 298 | 0.002 | 0.698 |
| **OrbN** | Bilateral orbitofrontal cortex; Bilateral hippocampus; Bilateral sup/mid occipital gyrus. | 295 | 0.003 | 0.681 |
| **SFN** | Bilateral superior/middle frontal gyrus; Bilateral angular gyrus; Bilateral precuneus; Right middle temporal gyrus. | 285 | 0.006 | 0.624 |
| **MTLN** | Bilateral hippocampus; Bilateral parahippocampus; Bilateral fusiform gyrus; Bilateral amygdala; Bilateral temporal pole; Bilateral middle temporal gyrus. | 270 | 0.019 | 0.538 |

Five of the identified networks were significantly disengaged (see Table 4) while the participants were listening to the musical excerpts: a *right Fronto-Parietal Network* (rFPN), covering right inferior and middle frontal, angular and inferior parietal gyri; a S*ensory-Motor Network* (SMN) comprising premotor and supplementary motor regions, the postcentral gyrus and basal ganglia; *Medial and Lateral Visual Networks* (MVISN and LVISN), extending mainly through the occipital lobe; and finally the *Default Mode Network* (DMN) comprised of its typical constituents, including the anterior and posterior cingulate cortex, the precuneus, and parietal regions.

**Table 4.** ICA networks *negatively* modulated by music listening. The main areas of the network are reported using a p < 0.05 FWE-corrected threshold at the cluster level. The level of network engagement with the task is reported using Wilcoxon signed-rank tests and FDR-corrected p-values. rFPN: right Fronto-Parietal Network; SMN: Sensorimotor Network; MVISN: Medial Visual Network; LVISN: Lateral Visual Network; DMN: Default Mode Network.

| **Network** | **Activation Region** | ***W*** | ***p<sub>FDR</sub>*** | ***r<sub>rb</sub>*** |
|---|---|---|---|---|
| **rFPN** | Right sup/inf frontal gyrus; Bilateral middle frontal gyrus; Right precentral gyrus; Right postcentral gyrus; Right insula; Bilateral ant/mid cingulate cortex; Right angular gyrus; Right supramarginal gyrus; Bilateral inferior parietal gyrus; Right mid/inf temporal gyrus; Right precuneus; Right mid/sup occipital gyrus; | 0 | p<0.001 | -1 |
| **SMN** | Bilateral precentral gyri; Bilateral postcentral gyri; Supplementary motor area; Bilateral middle cingulate gyri; Bilateral thalamus; Bilateral caudate; Bilateral putamen; Bilateral sup/inf parietal gyrus; Bilateral supramarginal gyri; Bilateral cuneus | 0 | p<0.001 | -1 |
| **MVISN** | Bilateral calcarine; Bilateral cuneus; Bilateral lingual gyrus; Bilateral precuneus; Bilateral middle/superior occipital gyrus; Bilateral precentral gyrus. | 17 | p<0.001 | -0.903 |
| **LVISN** | Bilateral sup/mid/inf occipital gyrus; Bilateral lingual gyrus; Bilateral posterior superior/mid/inf temporal gyrus; Bilateral fusiform gyrus; Bilateral inferior parietal gyrus; Bilateral postcentral gyrus; Bilateral precentral gyrus. | 26 | p<0.001 | -0.852 |
| **DMN** | Bilateral ant/post/mid cingulate cortex; Bilateral precuneus; Bilateral cuneus; Bilateral sup/mid occipital gyrus; Bilateral calcarine; Bilateral lingual gyrus; Bilateral angular gyrus; Bilateral inferior parietal gyrus; Bilateral sup/middle frontal gyrus; Thalamus; Bilateral middle temporal; Bilateral hippocampus; Bilateral parahippocampus; Bilateral fusiform gyrus; Brainstem. | 53 | p=0.002 | -0.698 |

#### ICA network modulation by overall and continuous liking ratings

When assessing which networks *were modulated by the liking judgments* made by the participants for each musical excerpt, we found significant results for both the overall and continuous ratings. Note that parametric beta values for high and low rated songs were corrected using the Medium trials as a baseline to account for individual differences in responses to aesthetic stimuli (corrected betas: High_m_ and Low_m_; see Materials and Methods).

The VN significantly responded to both High_m_ (W = 296, p_FDR_ = 0.021, r_rb_ = 0.68) and Low_m_ (W = 289, p_FDR_ = 0.031, r_rb_ = 0.64) overall liking judgments. No differences were found for the High_m_ > Low_m_ contrast for the VN (W = 178, p_FDR_ = 0.96, r_rb_ = 0.014; see Figure 2A), with this network being *equally modulated* by positive and negative overall judgments of liking (BF_01_ = 4.84). No other networks were significantly modulated by overall ratings or showed significant differences in engagement between the High_m_ and Low_m_ parametric modulators after correction for multiple comparisons.

**Figure 2.**
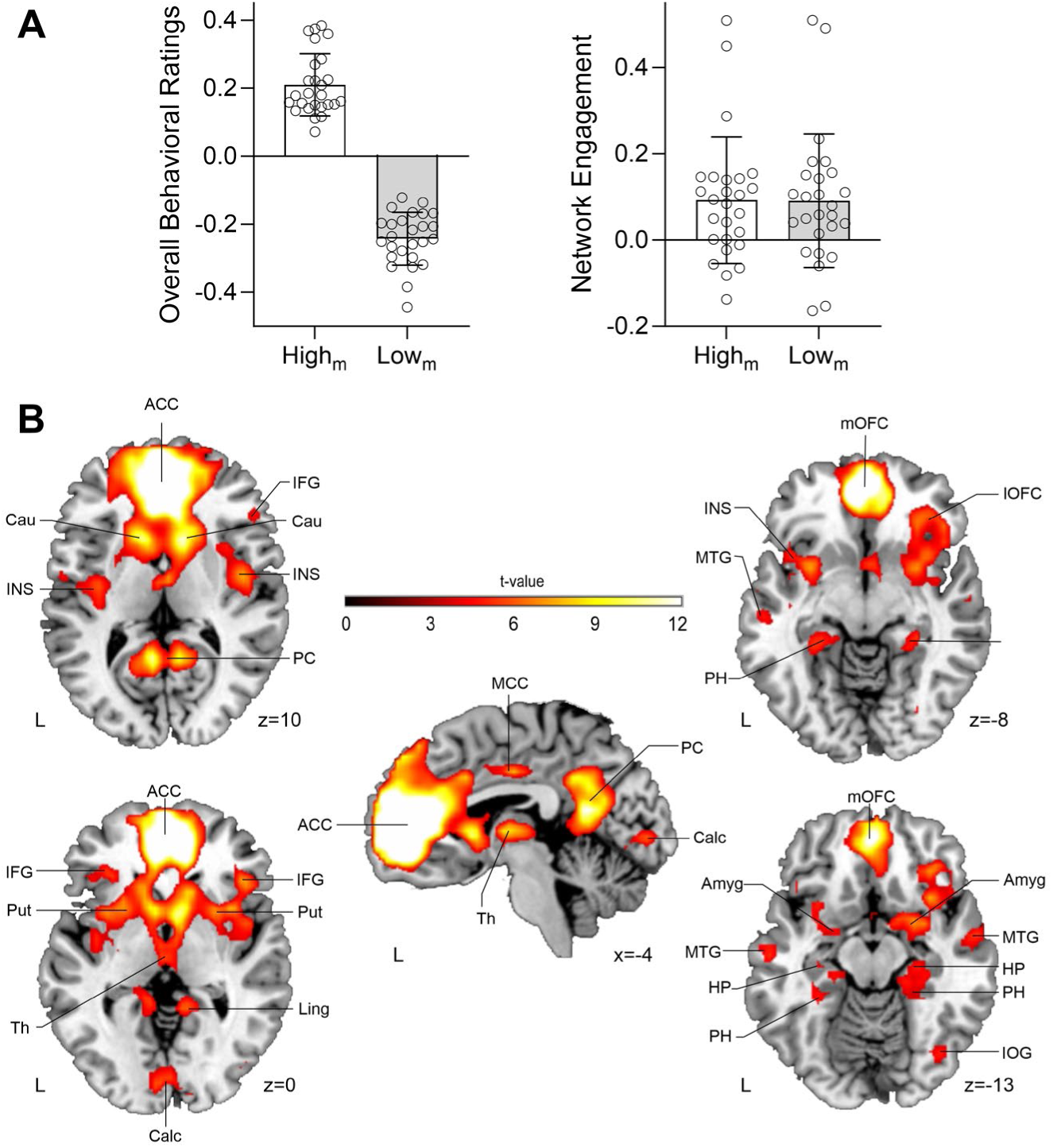
Network modulated by overall liking judgments. **A.** On the left, mean (plus SD) behavioral overall (i.e., discrete) liking ratings for High_m_ (white) and Low_m_ (grey) trials (corrected using the Medium behavioral ratings). This is the expected distribution of corrected ratings, since we classified songs into high or low conditions based on the overall ratings (we did not compute statistics to compare between overall High_m_ and Low_m_ overall ratings to avoid double-dipping). On the right panel, mean beta values (plus SD; corrected using Medium beta values), showing that the VN equally responds to High_m_ (white) and Low_m_ (grey) trials. Each circle represents data from one participant. This suggests that the VN is modulated by the overall aesthetic judgment of liking for both highly liked and disliked musical pieces. **B.** In red-yellow, spatial map corresponding to the VN (p < 0.05 FWE-corrected at the cluster level). Neurological convention is used with MNI coordinates at the bottom right of each slice. L, Left Hemisphere; ACC, Anterior Cingulate Cortex; MCC, Middle Cingulate Cortex; Cau, Caudate; IFG, Inferior Frontal Gyrus; INS, Insula; PC, Precuneus; Put, Putamen; Ling, Lingual Gyrus; Th, Thalamus; Calc, Calcarine; mOFC, medial Orbitofrontal Cortex; lOFC, lateral Orbitofrontal Cortex; MTG, Middle Temporal Gyrus; PH, Parahippocampal Gyrus; Amyg, Amygdala; HP, Hippocampus; IOG, Inferior Occipital Gyrus.

Since the VN resembles a network previously known to respond to stimuli with both positive and negative emotional valence, and in the presented data responds to both high and low overall liking ratings, we conducted an additional exploratory analysis. We compared the spatial component of the VN with several fMRI meta-analyses. First, we used NeuroSynth (a platform for large-scale, automated meta-analysis of fMRI data; www.neurosynth.org; [158]) to calculate an association map based on the search term “*valence”* that resulted in 361 studies (search performed on December 19^th^, 2024). Second, we obtained the frequency maps from a valence meta-analysis of 397 fMRI and PET studies for regions consistently activated by both positive and negative affective experiences [156]. As a control, we also included NeuroSynth association meta-analyses for *music* (163 studies; search performed on December 19^th^, 2024), *reward* (922 studies; search performed on December 19^th^, 2024), and a well-known meta-analysis of the DMN [141]. We then used the spatial sorting function provided by GIFT, which uses multiple linear regression to compare the spatial pattern of a particular ICA network with a set of templates [81]. Following this methodology, we spatially compared the VN with the meta-analyses of valence, reward, music, and the DMN. As a control, we repeated the same analysis, but using the DMN and APMN ICA networks obtained from our data (see Tables 3 and 4). We computed two multiple regressions, one per valence meta-analysis (i.e., one analysis included templates for the NeuroSynth meta-analysis of valence, reward, music, and the DMN from Smith and colleagues [141]; the other included templates for the meta-analysis of valence from Lindquist and colleagues [156], the NeuroSynth meta-analyses of reward and music, and the DMN from Smith and colleagues [141]). These analyses resulted in correlation coefficients (*r*) that indicated the degree of spatial similarity between each ICA network being analyzed (VN, DMN, APMN) and each template.

As expected, the highest *r* values for the APMN and DMN were those corresponding to the music and DMN meta-analyses, respectively. For the VN, the highest *r* corresponded to both meta-analyses related to *valence* (see Tables 5 and 6), and not to those corresponding to music, reward, or the DMN. This further suggests that the liking judgments provided by the participants—regardless of their *positive or negative value*—were supported by a series of brain regions that can respond to stimuli with *positive and negative emotional valence* rather than by a network more specifically related to reward [28,31] or music processing in general [159–161].

**Table 5.** Spatial sorting of ICA networks. The results shown here correspond to the multiple regression that included the **NeuroSynth valence meta-analysis**. *r* values show correlations between ICA networks and templates, with the highest correlation value for each ICA network shown in bold.

| **ICA Network** | **Valence**<br>NeuroSynth Meta-Analysis | **Music**<br>NeuroSynth Meta-Analysis | **Reward**<br>NeuroSynth Meta-Analysis | **DMN**<br>From Smith et al. [141] |
|---|---|---|---|---|
| **VN** | **0.201** | -0.016 | 0.050 | 0.061 |
| **DMN** | -0.014 | -0.008 | -0.0002 | **0.244** |
| **APMN** | -0.002 | **0.663** | -0.010 | 0.004 |

**Table 6.** Spatial sorting of ICA networks. The results shown here correspond to the multiple regression that included the valence meta-analysis from Lindquist and colleagues [156]. *r* values show correlations between ICA networks and templates, with the highest correlation value for each ICA network shown in bold.

| **ICA Network** | **Valence**<br>From Lindquist et al. [156] | **Music**<br>NeuroSynth Meta-Analysis | **Reward**<br>NeuroSynth Meta-Analysis | **DMN**<br>From Smith et al. [141] |
|---|---|---|---|---|
| **VN** | **0.351** | -0.012 | 0.052 | 0.061 |
| **DMN** | 0.033 | -0.007 | -0.002 | **0.244** |
| **APMN** | -0.017 | **0.663** | -0.01 | 0.004 |

Regarding continuous ratings, there was a significant difference between the engagement of the DMN by the slopes for the High_m_ > Low_m_ contrast (W = 62, p_FDR_ = 0.033, r_rb_ = −0.64; see Figure 3A and B). Specifically, for highly liked excerpts, more positive rating trends were associated with a more negative DMN response than for disliked excerpts. In other words, as ratings shifted upward, the DMN showed greater deactivation for music that listeners liked. No other networks were significantly modulated by the slope of the continuous ratings or showed significant differences in engagement between the High_m_ and Low_m_ parametric modulators after correction for multiple comparisons.

**Figure 3.**
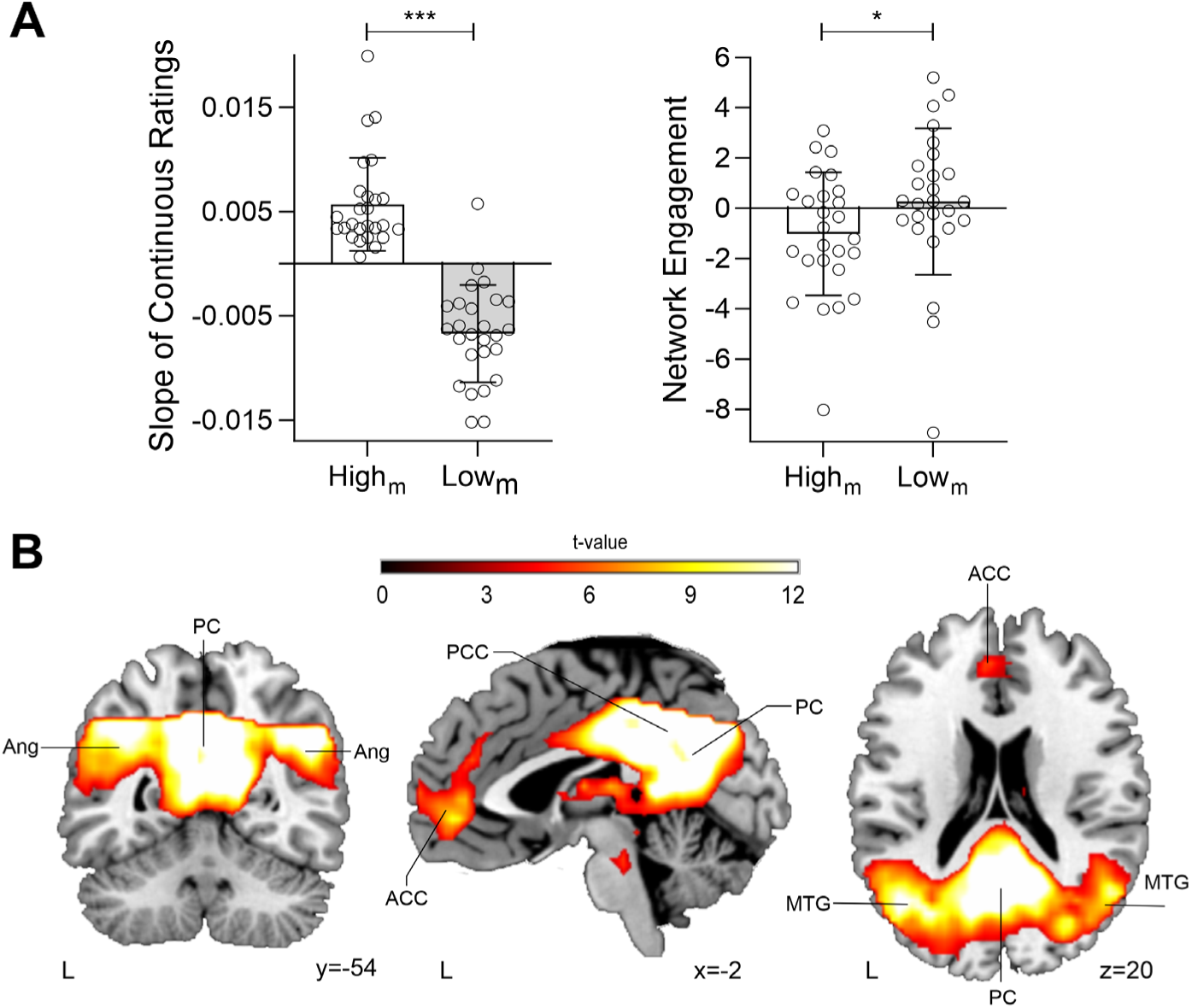
The Default Mode Network. **A.** On the left, mean (plus SD) behavioral slopes (summarizing the direction and average rate of change in continuous liking ratings across each excerpt) for High_m_ (white) and Low_m_ (grey) trials (corrected using the Medium behavioral slopes). In this case, since we classified songs into the highly liked or disliked condition using the overall ratings and not any measure extracted from the continuous ratings, we can compute statistics without double dipping. The corrected slopes were significantly higher for High_m_ than for Low_m_ (Wilcoxon signed-rank test, W = 351, p<0.001, r_rb_ = 1). This indicates more positive rating trends for highly liked than for disliked excerpts. On the right panel, mean beta values (plus SD; also corrected using Medium beta values), showing that the DMN is more negatively modulated by High_m_ (white) than by Low_m_ (grey) slopes. For highly liked excerpts, more positive rating trends were linked to greater DMN deactivation than for disliked excerpts. Each circle represents data from one participant. **B.** In red-yellow, spatial map corresponding to the DMN (p < 0.05, FWE-corrected at the cluster level). Neurological convention is used with MNI coordinates at the bottom right of each slice. L, Left Hemisphere; ACC, Anterior Cingulate Cortex; PCC, Posterior Cingulate Cortex; PC, Precuneus; MTG, Middle Temporal Gyrus; Ang, Angular Gyrus. * p < 0.05,*** p<0.001.

To further clarify the possible functional roles of the VN and DMN in aesthetic appreciation, we computed one final exploratory analysis focusing on individual differences in aesthetic responsivity. Using Spearman’s correlation, we assessed the relationship between the AReA [114] and the general engagement of the VN and DMN with the musical excerpts (using the beta values from the simplest model that only included one general musical condition). There was a significant correlation between the engagement of the VN and the AReA: the higher a participant’s general responsiveness to artistic stimuli, the more the VN was engaged while listening to music (r = 0.496, p = 0.01; see Figure 4). There was no significant relationship between the AReA and DMN (r = 0.331, p = 0.098).

**Figure 4.**
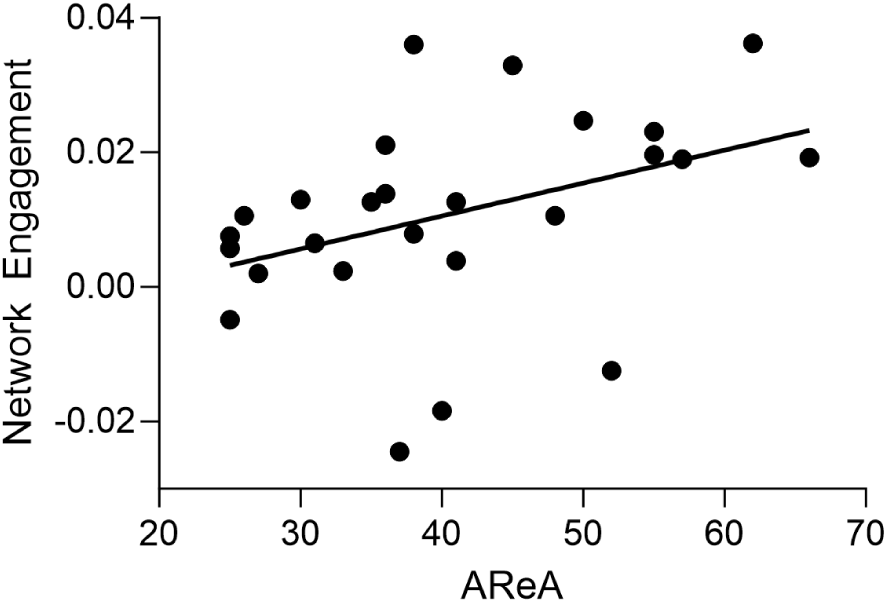
Individual differences in aesthetic responsiveness and VN engagement. The scatter plot shows the significant association (r = 0.496, p = 0.01) between the AReA (which measures individual differences in general responsiveness to artistic stimuli) and the network engagement of the VN with music (each dot represents data from one participant). As a measure of general network engagement to music, we used the beta values extracted from the simplest model that only included one general musical condition.

## DISCUSSION

This study assesses the neural underpinnings of both continuous and overall liking judgments of music. Behaviorally, as expected, there was low agreement in liking ratings across participants. Participants’ preferences were revealed very early in the presentation of a piece and were sustained over time [62,66], with the end value of the continuous liking judgement being the measure most highly related to the overall liking rating (see Figure 1). These behavioral results collected inside of the MRI scanner were replicated in a different sample of online participants (see Supplementary Figure 1). Regarding brain activity, the DMN showed more negative modulation by rating slope for highly liked than for disliked excerpts (Figure 3), a pattern that may reflect greater focused engagement with listening and evaluation as liking increases. Interestingly, a network comprised of regions known to respond to stimuli with both positive and negative emotional *valence* (the VN, see Figure 2, Table 3; [156]) was modulated by musical pieces rated both high and low on liking, with these liking ratings not depending on any acoustic parameters we measured. Our results regarding the VN extend to the individual differences domain, as participants with a greater general responsiveness to artistic stimuli engaged the VN more while listening to music (Figure 4).

One aim of this work was to investigate the relationship between continuous and overall ratings of liking during music listening. In agreement with previous research [123], we found that the final value (i.e., the value of the continuous trace at the end of an excerpt) showed the highest correlation with the overall ratings. While other metrics such as the average or peak value of the continuous ratings have also been related to overall ratings [73,123], in both our fMRI and online experiments the end value showed a stronger statistical relationship with the overall rating (see Figure 1C and Supplementary Figure 1C). This difference in the relationship of the average, peak, and final values with the overall rating could be due to the specific nature of the measures: here, listeners rated *liking*, while prior work often focused on ratings of *emotional intensity* of the music. It may also be the case that aesthetic judgments such as liking grow stronger over time, reflecting temporal integration [41,62,65,66]. As shown in Figure 1D and Supplementary Figure 1D, on average, liked pieces tended to be rated as increasingly more liked over the course of the trial, while ratings of disliked pieces grew more disliked. This suggests that listeners form a quick initial impression that strengthens (or is reinforced) during the course of the piece, which may explain why we found the strongest relationship between the end and overall ratings [72,162]. The results might also align with evidence accumulation models, which propose that humans make decisions by gradually gathering evidence in favor of a particular choice until a threshold is reached, triggering the decision [163,164]. Conversely, emotional intensity (e.g., chills) may show more variability over the course of the piece, which is why the peak or average may be a more representative metric of that reaction [30,32,68].

The fMRI analyses suggest an overlap between regions supporting liking judgments (the VN; see Figure 2) and areas previously known to be associated with the processing of emotional stimuli regardless of their positive or negative valence. Indeed, there was a spatial overlap between our VN and modality-general (i.e., not specific to musical stimuli; [165,166]) fMRI meta-analyses of valence (see Tables 5 and 6; [156,157,167]). Several of the regions that form the VN—including the amygdala, hippocampus, thalamus, striatum, and the ventromedial prefrontal cortex—have also been related to emotional processing of music and other stimuli [27–32,168]. While these studies usually focused on brain activity elicited during peak moments of high pleasure (e.g., chills), here we show that these regions are also modulated by the *liking* of the musical piece, both for high and low rated musical pieces.

It is noteworthy that individual differences in our participants’ preferences for abstract aesthetic stimuli (e.g., music, paintings, natural landscapes, architecture, dance, poetry) were also related to the engagement of the VN (see Figure 4), which suggests that this network might be related to modality-general aesthetic judgments [169]. This is in agreement with previous findings showing that in order to assign emotional value to abstract stimuli, an interaction between perceptual (e.g., visual, auditory) and reward, valuation and emotion-related circuits (e.g., the VN) is essential [36,66,170–173], and that individual differences in the structure and function of the neural substrates that support this interaction can affect the way we perceive and judge artistic stimuli [30,147,174–178]. Future research could examine whether activity in the VN can be used as a neural proxy for measuring individual differences in aesthetic judgments of liking across different domains (music, poetry, visual art, etc.).

The DMN showed more negative modulation by rating slope for highly liked than for disliked excerpts (Figure 3). One possible interpretation is that increasingly positive evaluations reflect greater engagement with the music and the listening task. Musical excerpts that listeners come to like more strongly may capture their attention, leaving fewer resources for unrelated, self-generated thoughts. The pattern may also reflect greater involvement in monitoring and updating their evaluations as the music unfolds. Both explanations are consistent with evidence linking stronger task-induced deactivation to increased processing demands and fewer task-unrelated thoughts [179,180]. Thus, the observed pattern may reflect a shift toward more focused listening and evaluation, although attention and thought content were not directly measured. However, recent accounts of the DMN offer an alternative interpretation by linking its activity to sense-making and the integration of new information into existing schemas [181]. As a well-liked musical excerpt unfolds, listeners may develop a clearer understanding of its structure, making subsequent musical events easier to integrate. The DMN effects could therefore reflect a diminishing need to revise their interpretation or update their schemas, even as listeners remain engaged.

Previous work in the visual domain has reported greater activation for high versus low appeal artworks in several regions of the DMN, including the medial prefrontal cortex and anterior cingulate, orbitofrontal cortex, inferior frontal gyrus and insula [92,101,182–185]. Engagement of the DMN has also been reported for highly aesthetically pleasing visual artwork [96,101,186,187]. However, those findings concern differences in aesthetic appeal, whereas the present DMN result concerns differences in how network activity relates to rating trends. These effects therefore need not be contradictory. One critical difference may be the role of prediction, expectations, and top-down processing for aesthetic experiences of visual art versus for extended musical pieces. Whereas a static image of a painting may invite active exploration, reflection, and top-down sense making, predictions and expectations in music are more directly determined by temporal, harmonic, and tonal structural properties of a composition that unfold over time and are therefore outside the listener’s control. During a musical clip, it is possible that both top-down predictions and aesthetic pleasure vary in a manner that leads to a less direct relationship between the overall aesthetic judgment at the end of the piece and average network engagement. In addition, it is possible that some musical clips are aesthetically pleasing for different reasons than visual art, such as the urge to dance (e.g., “groove” [188,189]) or to hum along, that rely less on top-down reflective processes thought to be mediated by the DMN. In this vein, differences in task demands may also contribute: evaluating music as it unfolds requires ongoing attention to a changing stimulus, whereas viewing a static artwork may offer different opportunities for reflection and self-referential processing.

The DMN has been shown to be also engaged during the aesthetic experience of music, with its activation influenced by stimuli of both positive and negative valence [11,17,100,190,191]. For example, previous research has shown that liked music resulted in higher engagement of DMN brain regions [95]. Other work showed that sad music (as compared to happy music) increased functional connectivity among regions within the DMN [94]. Moreover, one of the aforementioned studies showed that areas located in the DMN (e.g., precuneus) were activated for both liked and disliked music [95]. Here, during music listening, the DMN showed an overall negative response. For highly liked compared with disliked excerpts, more positive rating slopes were associated with a more negative DMN response (Figure 3). At first glance, this pattern may seem to contrast with previous research. However, the VN—which was influenced by both highly liked and disliked music—includes DMN regions such as the anterior cingulate cortex and precuneus (see Table 3). Moreover, as shown in Tables 5 and 6, the VN exhibits stronger spatial correlation with the DMN than with the broader meta-analysis of music, indicating an overlap between the VN and classical DMN areas. Because of this overlap and given that prior studies have reported increased DMN engagement during aesthetic experiences of music, our findings can be reconciled with those earlier observations. Another possible interpretation is that when regions of the DMN are engaged as parts of other networks (as happens here with the VN), overall DMN engagement may be suppressed; this can happen in some cases of music listening, especially as compared with engagement with visual art [192,193]. For example, listening to music or evaluating it may introduce processing demands that conflict with overall engagement of the DMN. By contrast, the DMN has been found to be engaged with listening to familiar music when individuals are not asked to make liking judgments (“free listening” paradigms; [95]), in line with task-specific suppression of the DMN. Finally, one possibility that could be explored in future studies is whether performing network decomposition on data collected during active music listening, vs. during independent rest scans, results in a different grouping of these overlapping regions.

One limitation of this study is that all participants had a predominantly Western musical background, which may limit the generalizability of our findings to other cultural contexts. In addition, we confined our study to two genres—classical and electronic—excluding “unconventional” compositions within those genres (e.g., atonal classical works) and omitting less popular styles such as punk or free jazz. Consequently, our results may not extend to situations where listeners, despite unfamiliarity or initial dislike of a genre, ultimately find themselves enjoying the music. In addition, participants gave higher average liking ratings to the classical excerpts than to the electronic excerpts. Thus, some of the behavioral and neural findings associated with highly liked vs disliked music may also partly reflect differences between the two genres.

In summary, we provide data linking behavior and brain function to better characterize a fundamental process: the way human listeners judge the aesthetic value of music. Participants determined whether they liked or disliked a particular piece (and maintained that decision) very early in time and their decisions—regardless of the positive or negative value of the ratings—were associated with a particular network composed of areas previously related to respond to modality-general valence. Our results extend to the individual differences domain, as we show that the engagement of the VN during music listening was associated with general aesthetic experience. Taken together, our results advance the hypothesis that our individual preferences in aesthetic choices are modulated by a complex interaction of circuits associated with emotion and reward.

## DATA AVAILABILITY

All behavioral data and ICA networks are provided as supplementary materials.

## AUTHOR CONTRIBUTIONS

Author contributions are listed following the CRediT (Contributor Roles Taxonomy) format. Pablo Ripollés: Data curation, Formal analysis, Investigation, Methodology, Validation, Visualization, Writing – original draft, Writing – review & editing. Amy M. Belfi: Conceptualization, Data curation, Formal analysis, Investigation, Methodology, Resources, Software, Writing – original draft, Writing – review & editing. Anna Kasdan: Data curation, Formal analysis, Investigation, Writing – review & editing. Edward A. Vessel: Conceptualization, Funding acquisition, Methodology, Project administration, Resources, Software, Supervision, Writing – review & editing. Andrea R. Halpern: Conceptualization, Methodology, Resources, Writing – review & editing. Jess Rowland: Conceptualization, Resources, Writing – review & editing. Robert Hopkins: Conceptualization, Writing – review & editing. G. Gabrielle Starr: Conceptualization, Funding acquisition, Project administration, Supervision, Writing – review & editing. David Poeppel: Conceptualization, Funding acquisition, Project administration, Supervision, Writing – review & editing.

## ACKNOWLEDGMENTS

We would like to thank Danielle Retcho for assistance with creating stimulus materials.

## SUPPLEMENTARY MATERIALS

**Supplementary Figure 1.**
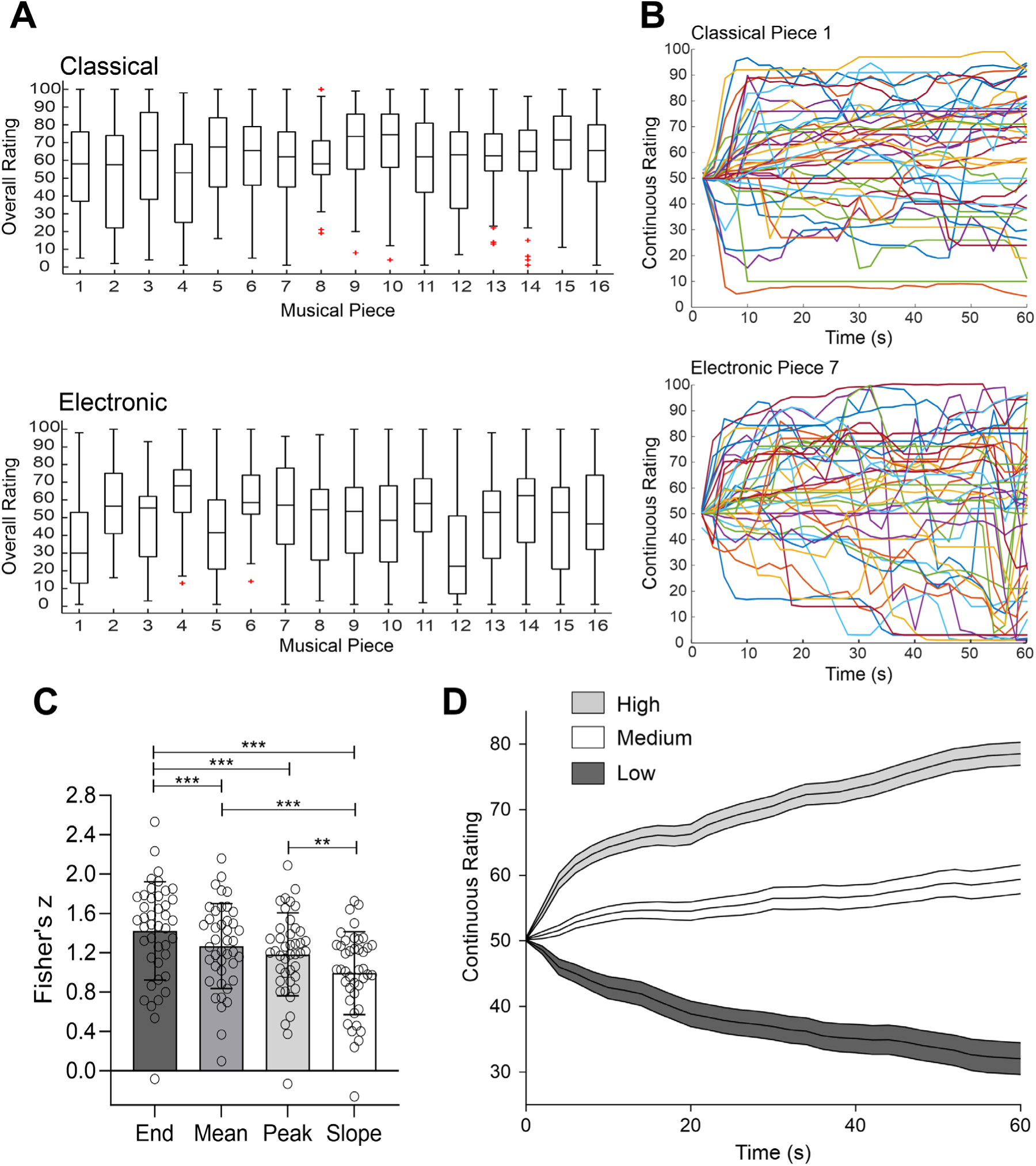
Online behavioral replication experiment. **A.** Boxplots illustrating participants’ overall ratings for each musical piece. Center line indicates median, bottom and top edges of the box indicate 25^th^ and 75^th^ percentiles. Whiskers extend to the most extreme data points not considered outliers, and red + symbols indicate individual outliers. **B.** Individual continuous responses from all participants for two representative musical excerpts (top, classical piece number 1; bottom, electronic piece number 7). Each line represents the continuous rating provided by a specific participant. **C**. Barplots (mean ± SD) showing the strength (using Fisher’s z transform) of the correlation between the *overall* rating and different measures extracted from the continuous ratings, including the *end, mean,* and *peak value*, and the *slope.* Each circle represents data from one participant. ** p < 0.005, *** p < 0.001. **D.** Mean (solid lines) plus SEM (shaded error bands) of *continuous* ratings (between 1 and 100) across participants are shown for High (grey), Medium (white) and Low (black) musical excerpts classified using the overall ratings.

